# Biomarker Candidates for Tumors Identified from Deep-Profiled Plasma Stem Predominantly from the Low Abundant Area

**DOI:** 10.1101/2021.10.05.463153

**Authors:** Marco Tognetti, Kamil Sklodowski, Sebastian Müller, Dominique Kamber, Jan Muntel, Roland Bruderer, Lukas Reiter

## Abstract

The plasma proteome has the potential to enable a holistic analysis of the health state of an individual. However, plasma biomarker discovery is difficult due to its high dynamic range and variability. Here, we present a novel automated analytical approach for deep plasma profiling and applied it to a 180-sample cohort of human plasma from lung, breast, colorectal, pancreatic, and prostate cancer.

Using a controlled quantitative experiment, we demonstrate a 257% increase in protein identification and a 263% increase in significantly differentially abundant proteins over neat plasma.

In the cohort, we identified 2,732 proteins. Using machine learning, we discovered biomarker candidates such as STAT3 in colorectal cancer and developed models that classify the disease state. For pancreatic cancer, a separation by stage was achieved.

Importantly, biomarker candidates came predominantly from the low abundance region, demonstrating the necessity to deeply profile because they would have been missed by shallow profiling.

## Introduction

Proteins control most biological processes in life. Alterations in their expression level, localization and proteoforms are often correlated with disease onset and progression^1^. In humans and animals, blood flows through virtually all tissues. Therefore, it has the potential to indicate the health state of any inner organ, even those not accessible from the outside. Blood is readily obtainable with minimal invasive sampling, and large biobanks exist for retrospective analyses^2^. Clinical analysis of blood is the most widespread diagnostic procedure in medicine, and blood biomarkers are used to diagnose diseases, categorize patients and support treatment decisions. Despite more than 20,000 diseases reported to affect humans^3^, it is only for a small fraction of them that accurate, sensitive and specific diagnostic tests exist.

The limited success of blood protein biomarkers is primarily due to analytical challenges that come with the proteomic analysis of blood plasma. On the one hand, the large biological variance between individuals and within individuals over time makes the discovery of reliable biomarker signatures difficult^4–7^. Further, the steep dynamic range of human plasma, with an estimated dynamic range of 12-13 orders of magnitude^8^, renders comprehensive proteome profiling challenging to any analytical technique. In the lower concentration range reside thousands of proteins, mostly tissue leakage proteins and signaling molecules that could serve as biomarkers but are very challenging to measure, especially in an unbiased manner^9,10^.

Mass spectrometry (MS)-based plasma analysis provides an unbiased, quantitative and therefore ideal technology for the system-wide characterization of the proteome^11^. Recently, technological developments in sample preparation, chromatography and acquisition enabled automated, large-scale plasma projects of hundreds of specimens that have resulted in reproducible findings^9,12–15^. These approaches share the shallow depth of proteome coverage, reaching a maximum of about 600 proteins identified and quantified in a sample. From qualitative analysis, disproportionately more proteins were found to be present in the lower abundance region of plasma than in the higher concentration range^10^. Novel MS-based approaches have been developed to improve analytical depth while retaining quantitative information. These include depletion of high-abundance proteins, enrichment of low abundant proteins of interest and prefractionation^16^. Still, they have yet to reach the throughput level needed to measure larger cohorts of clinical samples. Automatization and depletion-, batch- and quality-control have been tackeled^13,17,18^, but require further improvement for large scale studies. In summary, while current plasma proteome biomarker research approaches mostly cover the first few hundred proteins by concentration, rigorous experimental design and comprehensive, large-scale quantitative studies will achieve generalizable biomarker discovery^11^.

Screening for the most common cancer types cannot be done in a routine and population-wide manner. To date, only a few non-ideal, validated biomarkers exist in clinical use^19^. A significant challenge is that generally, only a single analyte or metric is measured despite the known heterogeneity of cancer. Biomarkers that accurately enable early detection in asymptotic subjects, reflect cancer aggressiveness at diagnosis and improve risk stratification are urgently needed^19^. Despite the medical need, plasma biomarker candidates for cancer are rarely validated or transferred to the clinic. Recent examples are: Zhang *et al.* performed discovery proteomics in plasma of 10 patients with colorectal cancer, discovered 72 biomarker candidates, and then did a successful follow-up verification for prognostic markers with 419 patients using an immunoassay^20,21^. Enroth *et al.* found plasma protein biomarker signatures for ovarian cancer^22^, but performed no validation. He *et al.* showed that for hepatocellular carcinoma and cholangiocarcinoma, biomarker candidates could be identified from plasma; validation of these candidates is still pending^23^. Zhou *et at.* identified biomarkers for early gastric cancer from a small sample set, but validation is still pending^24^. For prostate cancer, a blood diagnostic test was successfully developed based on discovery proteomics and is now being used in the clinic^25^. For detection of early ovarian cancer, the OVA1 test was developed and approved, where the measurement of beta-2 macroglobulin, apolipoprotein 1, serum transferrin and prealbumin is combined with the previously established marker CA125 to deliver better care^26,27^. This case exemplifies that multi-measurement techniques are expected to outperform single biomarker panels. Furthermore, single protein biomarkers are rarely specific for a single disease, e.g., Alpha fetoprotein is diagnostic in liver cancer, but the biomarker is not specific, as it is altered in other liver diseases, ovarian and testis cancer^28^. Rarely, there are highly specific biomarkers such as beta Subunit HCG (β-HCG), which is a serum marker for testicular carcinoma as β-HCG is never detected in the circulation of healthy men^29^. To make plasma biomarker discovery more efficient and successful, the comprehensive profiling and validation of large cohorts of plasma proteomes needs to be significantly improved with new approaches^11^. The expected outcome is new biomarkers that will allow early cancer detection and prediction of the probable response to therapy (in precision medicine).

We demonstrate a novel, automated analytical approach for plasma profiling to a depth of 2,732 proteins in the presented cancer study and identifying deep into tissue leakage and signaling molecule areas. We demonstrate identification and quantitative benefits over neat plasma profiling by a controlled quantitative experiment. Further, we profiled deep into the tissue leakage plasma samples coming from both healthy patients and patients with one of the five most deadly solid tumors in the United States^30^. A biomarker analysis with machine learning revealed candidates and models able to classify healthy and diseased samples. The discovered biomarker candidates predominantly came from low abundance protein regions, clearly demonstrating the need to measure deeply because they would have been missed by shallow plasma profiling.

## Experimental Procedures

### Ethics

The Cantonal Ethics Committee for Research on Human Beings, Zürich, Switzerland approved the study protocol to be performed (Proteomic analysis of plasma samples (2020-02892)).

### Cohort selection and study design

Cohort selection and experimental design was driven by sample availability in commercial repositories. For each cancer type, 30 matching samples were selected and split into early (non-metastatic stage IA-IIC) and late (non-metastatic stage IIIA-C) groups. Prior to the analysis, normal individuals were matched for age, sex and whenever possible balanced across ethnicities to both early and late groups for each cancer type. This resulted in three equal control groups (n=15) with overlapping individuals, namely: breast cancer control, prostate cancer control and remaining cancer control. Matching was done manually using the χ2 test or ANOVA with a p-value threshold at 0.05 (R-package ‘tableone’).

### Sample preparation of the pan-cancer cohort

180 human plasma samples were obtained from Precision for Medicine and its subsidiaries (Norton USA), Discovery Life Sciences (Huntsville, USA) and ProteoGenex (Los Angeles, USA). Due to limited availability, samples were not balanced across suppliers; collection procedures and handling until storage at −80°C are considered to be the same in the case of all three providers (Supplementary Table 1). All samples were handled equally and thawed twice. During the aliquoting, a small amount of each sample was pooled. This quality control sample was subsequently used for the library generation and to assess quality and batch effects throughout the sample preparation and acquisition. The processing batches were block randomized for disease status, disease state, gender and ethnicity (only relevant for breast cancer samples) and kept for the entire sample preparation.

Depletion was performed using the Agilent Multi Affinity Removal Column Human-14, 4.6 × 50 mm (Agilent Technologies) set up on a Dionex Ultimate 3000 RS pump (Thermo Fisher Scientific) and run according to the manufacturer’s instructions. Briefly, the plasma was diluted 4:1 with Buffer A for Multiple Affinity Removal LC Columns (Agilent Technologies) and filtered through a 0.22 μm hydrophilic PVDF membrane filter plate (Millipore) before 70 μl were injected onto the column. The gradient was 27.5 min long, with the collection occurring between 3.6 and 9.2 min, a flow rate of 1 ml/min during 11 and 26.5 min and 0.125 ml/min during the rest of the gradient, and Buffer B for Multiple Affinity Removal LC Columns (Agilent Technologies) only in the time period 13 to 17.5 min (100% Buffer B). Well-spaced within each processing batch, we depleted the quality control sample three times and treated it as a separate sample thereon (depletion control samples).

Following depletion, we digested the samples with protein aggregation capture using a KingFisher Flex (Thermo Fisher Scientific)^31^. To assess digestion reproducibility, we mixed two extra depletions of the quality control sample before splitting it into digestion triplicates (digestion control samples). The acidified peptide mixtures were loaded for cleanup into MacroSpin C18 96-well plates (The Nest Group), desalted, and eluted with 50% acetonitrile. Samples were dried in a vacuum centrifuge, solubilized in 0.1% formic acid, 1% acetonitrile with Biognosys’s iRT and PQ500 kits (Biognosys) spiked following the manufacturer’s instruction. Prior to DIA mass spectrometric analyses, the sample’s peptide concentrations were determined using a UV/VIS Spectrometer at 280 nm/430 nm (SPECTROstar Nano, BMG Labtech) and centrifuged at 14,000 × g at 4 °C for 30 min.

### Sample preparation of the Controlled Quantitative Experiment

The controlled quantitative experiment was generated from 20 healthy human EDTA K3 plasma samples obtained from Sera Laboratories International Ltd. (West Sussex, UK). *Saccharomyces cerevisiae* (*S*. *cerevisiae*) were lysed in 100 mM HEPES pH 7.4, 150 mM KCl, 1 mM MgCl_2_, by shear force passing through a gauge 12 syringe for 15 times on ice before filtering (0.2 μm). *Escherichia coli (E. coli)* was lysed with a cell cracker before filtering (0.2 μm). After protein concentration determination using a UV/VIS Spectrometer at 280 nm (SPECTROstar Nano, BMG Labtech), each sample was spiked with fixed ratios of *E*. *coli* and *S*. *cerevisiae* leading to a synthetic 1:2 and 4:3-fold change. To 20 μl plasma (∼1200 μg proteins), 40 or 30 μg *S*. *cerevisiae* and 12 or 24 μg *E*. *coli* lysate were added for condition A and B, respectively. The resulting 40 samples were diluted 4:1 with Buffer A for Multiple Affinity Removal LC Columns (Agilent Technologies), filtered through a 0.22 μm hydrophilic PVDF membrane filter plate (Millipore). 70 μl were used for depletion as described above followed by Filter-Aided Sample Preparation (FASP)^32^ and 30 ul for the neat plasma comparison. The diluted neat plasma sample was precipitated by adding four excesses of cold acetone (v/v) and overnight incubation at −20 °C. The pellet was subsequently washed twice with cold 80% acetone in water (v/v). After air-drying the pellet, the proteins were resuspended in 50 μl denaturation buffer (8 M Urea, 20 mM TCEP, 40 mM CAA, 0.1 M ABC), sonicated 5 minutes (Bioruptor plus, Diagenode, 5 cycles high, 30 s on, 30 s off) and incubated at 37 °C for 60 min. Upon dilution with 0.1 M ABC to a final urea concentration of 1.4 M, the samples were digested overnight with 2 μg sequencing grade trypsin (Promega) and trypsin inactivated by adding TFA to a final concentration of 1% v/v. Peptide clean-up was carried out as described above.

### Library generation

High pH reverse phase (HPRP) fractionation was performed using a Dionex UltiMate 3,000 RS pump (Thermo Fisher Scientific) on an Acquity UPLC CSH C18 1.7 μm, 2.1×150 mm column (Waters) at 60 °C with 0.3 ml/min flow rate. Prior to loading, the pH of 300 μg of pooled samples was adjusted to pH 10 by adding ammonium hydroxide. The used gradient was 1% to 40% solvent B in 30 minutes; solvents were A: 20 mM ammonium formate in water, B: acetonitrile. Fractions were taken every 30 seconds, sequentially pooled to 20 fraction pools. The fraction pools were then dried down and resuspended in 0.1% formic acid, 1% acetonitrile with Biognosys’s iRT kits spiked in according to the manufacturer’s instruction. Before DDA mass spectrometric analyses, peptide concentrations were determined, and the samples were centrifuged as described above.

### Mass spectrometric acquisition

For DIA LC-MS measurements, 1 μg of peptides per sample was injected onto an in-house packed reversed-phase column (PicoFrit emitter with 75 μm inner diameter, 60 cm length and 10 μm tip from New Objective, packed the Reprosil Saphir C18 1.5 μm phase (Dr. Maisch, Ammerbuch, Germany) on a Thermo Fisher Scientific EASY-nLC™ 1,200 nano-liquid chromatography system connected to a Thermo Fisher Scientific Orbitrap Exploris 480 mass spectrometer equipped with a Nanospray Flex™ ion source. The DIA method was adopted from Bruderer et al.^33^ and consisted of one full-range MS1 scan and 29 DIA segments.

For DDA and DIA LC-FAIMS-MS/MS measurements, 4 μg of each sample was separated using a self-packed analytical PicoFrit column (75 μm × 50 cm length) (New Objective, Woburn, MA, USA) packed with ReproSil- Saphir C18 1.5 μm (Dr. Maisch GmbH, Ammerbuch, Germany) with a 2 hours segmented gradient using an EASY-nLC 1,200 (Thermo Fisher Scientific). LC solvents were A: water with 0.1 % FA; B: 20 % water in acetonitrile with 0.1 % FA. For the 2 hours gradient, a nonlinear LC gradient was 1 - 59 % solvent B in 120 minutes followed by 59 - 90 % B in 10 seconds, 90 % B for 8 minutes, 90 % - 1 % B in 10 seconds and 1 % B for 5 minutes at 60°C and a flow rate of 250 nl/min. The samples were acquired on an Orbitrap Exploris 480 mass spectrometer (Thermo Fisher Scientific) equipped with a FAIMS Pro device (Thermo Fisher Scientific) using methods based on^34^. If not specified differently, the FAIMS-DIA method contained three FAIMS CVs (−35V, −55V, and −75V) parts with each a survey scan of 120,000 resolution with 20ms max IT and AGC of 3*10^6^ and 35 DIA segments of 15,000 resolution with IT set to auto and AGC set to custom 1,000%. The mass range was set to 350-1,650 m/z, the default charge state to 3, loop count to 1 and normalized collision energy to 30. For the acquisition of the fractionated sample for the library, a DDA method was applied. The DDA method consisted of three FAIMS CVs (−35V, −55V, and −75V): each contained a DDA experiment with 60,000 resolution of MS1, 15,000 resolution of MS2, with fixed cycle time (1.3s), IT set to AUTO and AGC set to custom 500%^35^.

### Mass spectrometric data analysis

#### Database Search for library generation

DIA and DDA mass spectrometric data were analyzed using the software SpectroMine (version 3.0.2101115.47784, Biognosys) using the default settings, including a 1% false discovery rate control at PSM, peptide and protein level, allowing for 2 missed cleavages and variable modifications (N-term acetylation and methionine oxidation). The human UniProt .fasta database (*Homo sapiens*, 2020-07-01, 20,368 entries) was used and for the library generation, the default settings were used except for the use of a top 300 precursors per protein filter.

#### Quantitative analysis of data independent acquisition

Raw mass spectrometric data were first converted using the HTRMS Converter (version 14.3.200701.47784, Biognosys) and then analyzed using the software Spectronaut (version 15.0.210108, Biognosys) with the default settings, but Qvalue sparse filtering was enabled with a global imputing strategy and a hybrid library comprising all DIA and DDA runs conducted in this study^36^. Default settings include peptide and protein level false discovery rate control at 1% and cross-run normalization using global normalization on the median. Including a high number of quality control samples (depletion, digestion and injection controls) enabled the investigation for batch effects and quantification of introduced variability at each step. No batch effect was identified by either principal component analysis (PCA, ‘stats’ R-package) or hierarchical clustering.

CQE DIA data were analyzed using the directDIA approach of Spectronaut software (version 15.0.210108, Biognosys) using the default settings, including a 1% false discovery rate control at PSM, peptide and protein level, allowing for 2 missed cleavages and variable modifications (N-term acetylation and methionine oxidation). The combined human, *E*. *coli* and *S*. *cerevisiae* .fasta databases with the removal of the overlapping tryptic sequences (*Homo sapiens* 2020-08-31, 96,996 entries; *Saccharomyces cerevisiae* (strain ATCC 204508 / S288c), 6,078 entries; *Escherichia coli* (strain K12), 4,857 entries; *Combined*, 96,637 entries) was used and for the library generation the default settings were used except for Qvalue sparse filtering enabled with a global imputing strategy and cross run normalization using global normalization on the median based solely on the human identifications.

When we use proteins, we refer to protein groups as determined by the ID picker algorithm^37^ and implemented in Spectronaut.

### Data analysis and biomarker selection

Initial univariate candidate filtering was performed using pairwise Wilcoxon test applied per protein across disease status (healthy, early and late stage) with Holmes-Bonferroni correction (within-group). Proteins with a p-value below or equal 0.05 from randomly selected 80% of observations were used for further optimization using sparse partial least square discriminant analysis (sPLSDA)^38^. A leave-one-out algorithm was used for optimal component and protein selection. sPLSDA training and testing were performed using the R-package ‘mixOmics’^39^. The remaining 20% of observations were used for validation. Accuracy of prediction for all three groups, healthy, early stages, late stage, and healthy against early and late stages together, were calculated as the ratio of the true positive and negative-sum to all observations (R-package ‘caret’). Unsupervised hierarchical analysis was done with Manhattan distance and Ward’s clustering on centered and normalized data (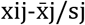, i-th observation with j-th protein) using R-package ‘ComplexHeatmap’. PCA analysis was done using R-package ‘stats’. Correlation analysis was done using Pearson correlation with R-packages ‘stats’ and ‘corrplot’. Correlation significance was tested using a two-sided t-test at 0.05 alpha. All analyses were performed using log_2_ transformed data. Gene ontology enrichment was performed using GOrilla^40^, the identifications of this study were selected as background. All basic calculations and data transformations were performed in R with R-packages: ‘dplyr’ and ‘ggplot2’.

## Results

### Optimization and validation of the analytical approach

While methods to analyze the plasma proteome in-depth exist, they are usually either targeted and therefore biased, as for the case of antibody or aptamer-based technologies, or are based on the principle of fractionation and are therefore difficult to scale. We aimed to develop an analytical method that provided deep coverage and quantitative accuracy while minimizing sample handling, bias and batch effects. For this scope, we developed and optimized an automated plasma depletion pipeline composed of three major steps: sequential depletion, parallel digestion and LC-MS acquisition (Fig. 1A).

**Fig. 1:**
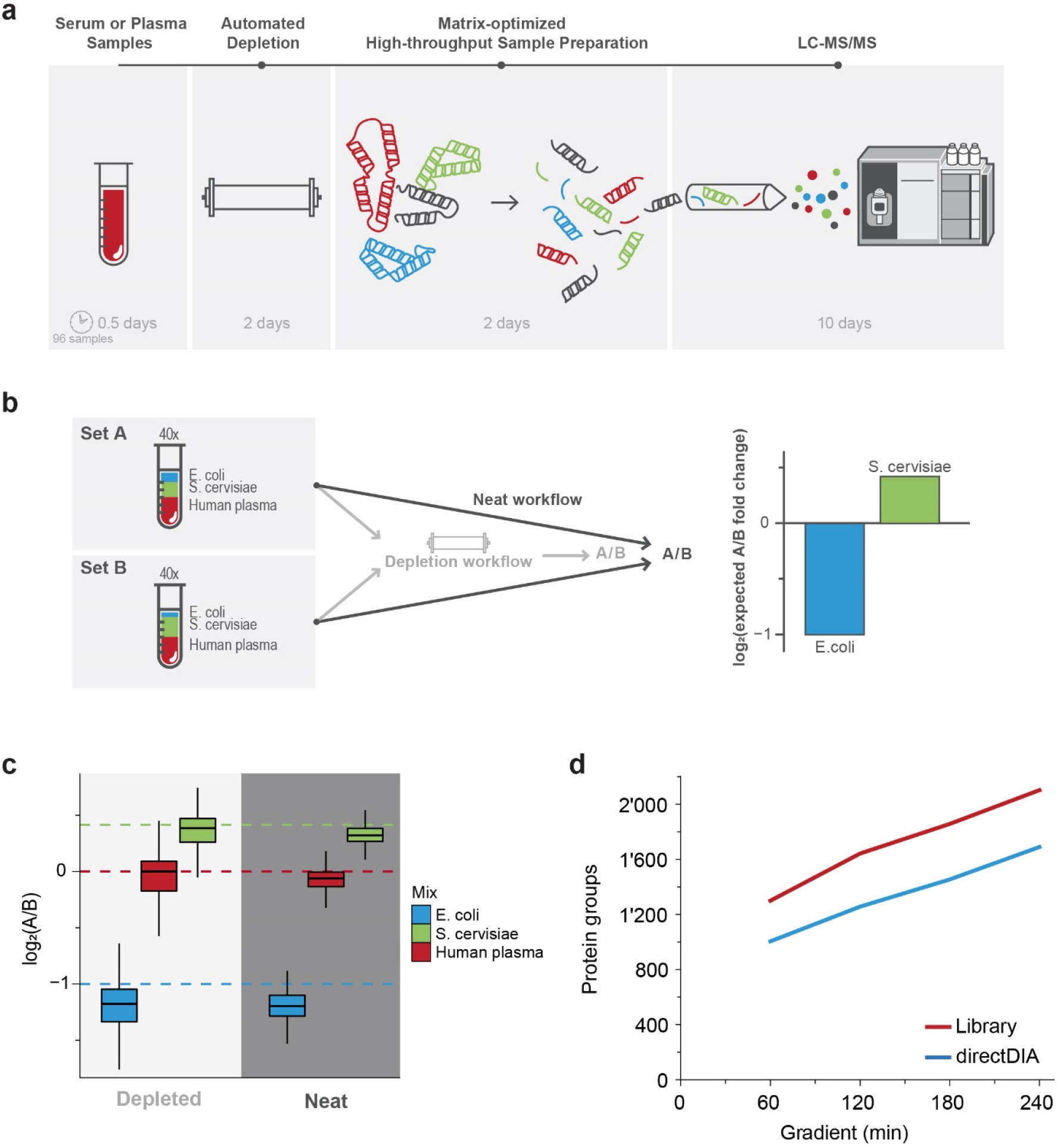
Deep plasma profiling: automated analytical approach and benchmarking. **(a)** Sketch of the major steps of the analytical approach developed for deep human plasma profiling for biomarker discovery, including depletion of the 14 most abundant proteins and the approximate time requirements. **(b)** Schema of the controlled quantitative experiment based on human plasma spiked with known amounts of *Saccharomyces cerevisiae (S. cerevisiae)* (1:1.3) and *Escherichia coli (E. coli)*. (1:.5). The controlled mixtures were either directly digested or processed using the process described in panel *a*. **(c)** Plot showing the measured distributions of the fold changes of the controlled quantitative experiment divided by species. The dashed lines represent the theoretical fold change. **(d)** Comparison of the number of protein groups identified at different gradient lengths for a depleted human plasma pool by either directDIA (blue) or with a sample specific library (red).

First, we automated the depletion of the 14 most abundant proteins using a sequential approach supporting a 96-well format^41^. Briefly, after randomization and filtration of the samples into a 96-well plate, an automated chromatographic system sequentially and automatically processed the plate, thereby depleting the 14 most abundant human proteins in plasma via the use of specific antibodies.

In order to quantify the analytical gain of the approach and to assess whether depletion maintains quantitative precision and accuracy, we performed a controlled quantitative experiment (CQE). The CQE sample set was generated from 20 healthy human plasma samples spiked with either 1:400 *E*. *coli* and 1:90 *S*. *cerevisiae* for condition A or 1:200 *E*. *coli* and 1:120 *S*. *cerevisiae* for condition B (Fig. 1B). After processing the 40 samples with or without the automated depletion pipeline, they were analyzed on a mass spectrometer using data independent analysis (DIA). Since the major challenge linked to quantification in plasma is the large dynamic range, removing the 14 most abundant proteins should lead to an increase in the number of proteins identified compared to the neat plasma. Indeed, while the processing of the neat plasma samples led to an average identification of 572 proteins (3,920 peptides) across all samples, depletion significantly increased coverage by 257% to 1,471 proteins (10,230 peptides) (n = 40, p-value = 1e-98, Supplementary Fig. 1A). Importantly, depletion retained the quantitative accuracy close to the expected ratios between condition B and A of 0.415 for *E*. *coli* and −1 for *S*. *cerevisiae*: *E*. *coli* median ratio −1.20 and −1.18 and *S*. *cerevisiae* 0.38 and 0.32 for the neat and depleted set, respectively (Fig. 1C). Finally, we performed an unpaired t-test between conditions B and A and could identify 171 and 621 candidates (FDR, q-value >= 0.01) for the neat and depleted set, respectively (Supplementary Fig. 1B). Given the experiment’s controlled nature, we could identify the true hits as those proteins mapping to either *E*. *coli* or *S*. *cerevisiae* and showing the expected directionality. Overall, depletion led to a 362% increase in true hits, 170 and 615 for neat and depleted (actual FDR < 1% for both), respectively. In summary, the automated depletion more than tripled the number of proteins identified and the number of true hits while maintaining quantitative accuracy and reducing the manual workload to only the filtering of the samples (about half a day per 96 samples, Fig. 1A).

In the second step following depletion, the sample plate was prepared for digestion on an automated platform using a protein aggregation capture approach^31^. Subsequently, the samples were cleaned using C18 plates, and peptide concentration was measured. In case a library was generated, a fraction of all samples can be pooled and an ultra high-pressure liquid chromatography controlled high pH reverse phase (HPRP) fractionation was performed^33^.

The third step comprises the LC-MS measurement of the samples. Even after depletion of the most abundant proteins, the major challenge hindering quantification is the large dynamic range in plasma. Hence, we developed and optimized the LC-MS acquisition for deep proteome coverage by using FAIMS-based ion mobility on the orbitrap platform combined with high-performance chromatography. We developed FAIMS-DIA methods that maximize protein and peptide identification by comparing values and counts of FAIMS compensation voltages with different scan resolutions. This resulted in a set of optimized methods for gradients from one to four hours. Benchmarking with the depleted plasma resulted in 1,300 protein identifications in one-hour gradients to 2,103 protein identifications in four hours (Fig. 1D). For reference, in the human cell line HeLa, 10,026 proteins were identified in four hours (Supplementary Fig. 1C).

Altogether, we demonstrated that the presented automated plasma depletion pipeline has the potential to enable the unbiased, reproducible and precise quantification of more than 2,000 proteins on average per sample across very large cohorts.

### How deep and accurately in the plasma proteome can we see

To test our pipeline, we set out to analyze a diverse cohort of human plasma samples coming from the five most deadly solid cancer types in the United States^30^: pancreatic, colorectal, breast, prostate and non-small cell lung cancer. For each cancer type 15 early (stage I to IIC) and 15 late stage (IIIA to IIIC) non-metastatic patients, as well as 15 matching normal control samples, were selected, based on available baseline data (including gender, age and where applicable smoking status, Fig. 2A and Supplementary Table 2). Altogether, we processed 180 samples (and an additional 24 quality control samples) over the course of one week and approximately a month of measurement time. With this scalable approach, we could identify and quantify 2,732 proteins (2,463 proteins with two or more peptide sequences) across 226 measurements (180 samples and 46 quality control samples, about 900 proteins/hour measurement, Fig. 2B), of which 1,804 are found in at least 50% of the runs (Supplementary Fig. 2A). With the identified proteins, we could cover the eight orders of magnitude dynamic range reported for plasma in the Human Protein Atlas (3,222 proteins detected in human plasma by mass spectrometry, of which we could quantify 70%, Supplementary Fig. 2B). Within this range, we extensively covered the tissue leakage proteome, interleukins and signaling proteins such as EGF, KLK3 (PSA), AKT1, CD86, MET, ERBB2 and CD33 (Fig. 2C). As expected, among the 500 highest intensity proteins, meaning the proteins that would likely be identified, if no depletion would have been applied, 196 (39%) are classified as secreted proteins. On the lower end, we identified tissue-specific proteins coming from the diseased organs (n = 42, 81% of which are not part of the 500 most abundant proteins), cytokines (n = 29, 85%) and nucleoplasm (n = 637, 90%) proteins exemplifying the different functional plasma concentration ranges (Fig. 2C). We identified 190 targets for FDA-approved drugs, of which 125 (66%) fall in the lower intensity range^42^. The different biological role of low and high abundant plasma proteins shows that we could recover the known biology of the plasma proteome.

**Fig. 2.**
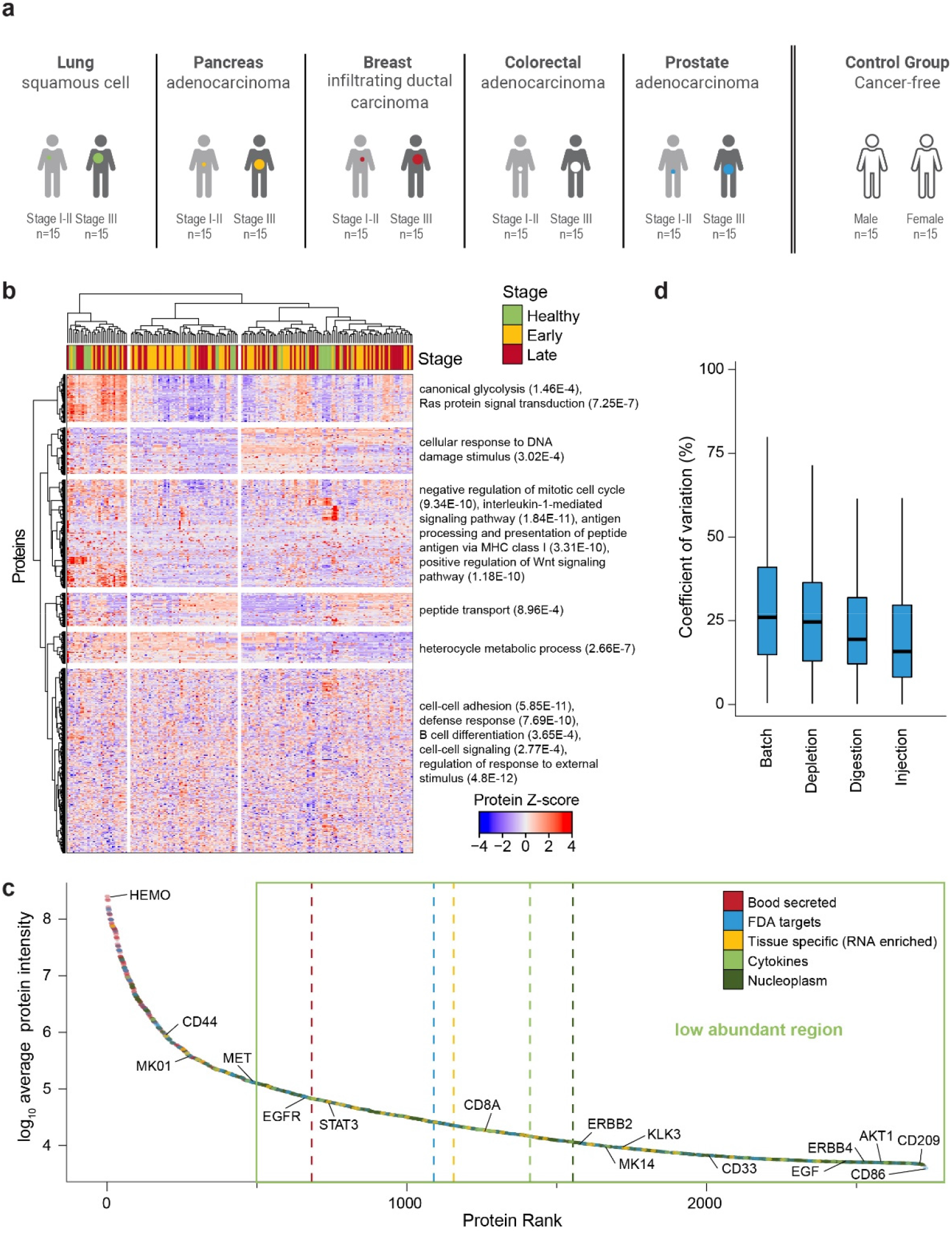
Deep plasma discovery proteomics of five solid cancer types. **(a)** Description of cohort comprising five solid cancers: breast (infiltrating ductal carcinoma), colon (adenocarcinoma), pancreas (adenocarcinoma), prostate (adenocarcinoma) and lung (non-small lung cancer, squamous cell) cancer. 15 subjects for early and late stages were selected for each cancer type, along with 15 matching healthy individuals (a total of 30, given the need to balance ethnicity and sex for prostate and breast cancer). **(b)** Z-score of all quantified proteins (n=2,732) across all measured samples (n=180). Stage calling is overlaid. Both the proteins and the samples were hierarchically clustered. Selected, significantly enriched gene ontology pathways are reported on the right with the p-value in brackets. **(c)** The protein rank vs. protein average intensity (n=180). Proteins were categorized according to Human Protein Atlas and the average rank was calculated (dotted, vertical lines). The green box depicts the proteome region that is typically below the sensitivity of neat plasma profiling by mass spectrometry. **(d)** The coefficient of variation (CV) of the quality control measurements across the processing steps was plotted. Controlled were LC-MS variance by reinjection of the same digested sample (injection), digestion and depletion were done repeatedly of the same sample (digest, depletion) and the batch stemming from sample preparation 96-well plates (batch). Thick lines indicate medians, boxes indicate the 25% and 75% quartiles, and whiskers extend between the median and ± (1.58 × interquartile range).

Furthermore, based on quality control samples, we could characterize variance introduced on each level: injection (median coefficient of variation (CV = 16%), digestion (CV = 19%), depletion (CV = 25%), and column (CV = 26%), all of which are much lower than the healthy inter-individual variability (CV = 56%, Fig. 2D and Supplementary Fig. 2C). As a further quality control, we focused on known protein levels’ inter-patient variability (measured by CV, Supplementary Fig. 2D). On one hand, coagulation- and complement cascade proteins (KEGG complement and coagulation cascades) were significantly enriched amongst the proteins with the least inter-patient variability, (median CV = 32% and p-value = 2.8e-12), such as complement factor I (CF1, CV = 23%) and complement component C6 (CV = 27%), demonstrating tight regulation^13^. On the other hand, keratins (likely contaminants, Go biological process keratinization) were significantly enriched amongst the proteins with the most inter-patient variability (CV = 339% and p-value = 4.46e-8), with HLA molecules (CV = 90%) also showing high variability across patients^43^. Additionally, lipoprotein A (LPA) showcases a large inter-patient variability (CV = 113%), likely due to the known genetic variants affecting its secretion into plasma^44,45^. Overall, the quantitative dataset generated recapitulates known biological features of intra-patient heterogeneity while providing a deep unbiased view of the plasma proteome.

### Considerable heterogeneity across cancer types

The cohort was designed to enable five independent within-cancer analyses, each comprising a healthy, an early and a late stage group (each n = 15, Supplementary Table 2, Fig. 2A). Overall, we included 30 control samples, but only a subset of 15 per cancer were matched (see methods). Hence, a combined analysis of all samples together was not the primary goal of this study. Aware of these limitations, we explored the entire dataset for markers that would agnostically predict the cancer stage. The analysis pipeline applied to the whole data set and the cancer-specific analyses were the same and aimed at providing actionable insights about specific disease development. Given the large amount of data (2,732 proteins combined), we performed a two-step approach (Fig. 3A). First, we filtered for differentially abundant proteins between healthy, early and late stage cancer using univariate analysis. In the case of the pan-cancer model, we found 468 proteins dysregulated (Fig. 3B, Supplementary Fig. 3A and Supplementary Table 3). Second, using the selected proteins, we trained a model based on sparse partial least square discriminant analysis (sPLSDA) on 80% of the data set. This modeling step further reduced the number of proteins to 94 (Fig. 3B). The model partially differentiated healthy from disease but not late to early stage (Supplementary Fig. 3B and Supplementary Table 4). Interestingly, the majority of the differentiating proteins would have been below the detection level in a neat plasma preparation (65%, Fig. 3C). Furthermore, the unsupervised clustering of the differentiating proteins generated enriched patterns (Fig. 3D). For example, proteins enriched for immunoglobulin production and complement activation tend to be higher in healthy samples (Fig. 3E). A subset of cancer samples has a strong upregulation of proteins linked to metabolic processes and cellular oxidant detoxification (Fig. 3D and E). Immunoglobulin kappa variable 6-21 (KV621) was among the proteins higher in healthy samples, was the third most important discriminant protein in the model (0.56 importance^46^), showed a more pronounced bi-modal distribution in healthy individuals, and a decrease in diseased individuals (Fig. 3F and Supplementary Fig. 3C). In addition, the model identified the known inflammation marker Complement C5 (CO5, importance 1) as increased in early and late stage and Spondin-1 (SPON1, importance 0.58) increased in late stage (Fig. 3F and Supplementary Fig. 3C), as the first and second most important contributors, respectively. Finally, the predictive power of the model was validated using the remaining 20% of the samples. The predictive power was low with 55.6% (Supplementary Fig. 3D), likely due to the cohort imbalance, the sample heterogeneity and the small sample set, as each cancer type is known to have a particular protein signature^47^. Nonetheless, unsupervised clustering using the final protein panel (enrichment p-value = 1.4e-9) allowed for more efficient separation of samples between healthy and disease states compared to the entire proteome (p-value = 0.09, Fig. 2B and 3D). Altogether, global data analysis underlined the importance and necessity of precision medicine and a much larger sample set would be needed to find a potential “one-fits-all” solution.

**Fig.3.**
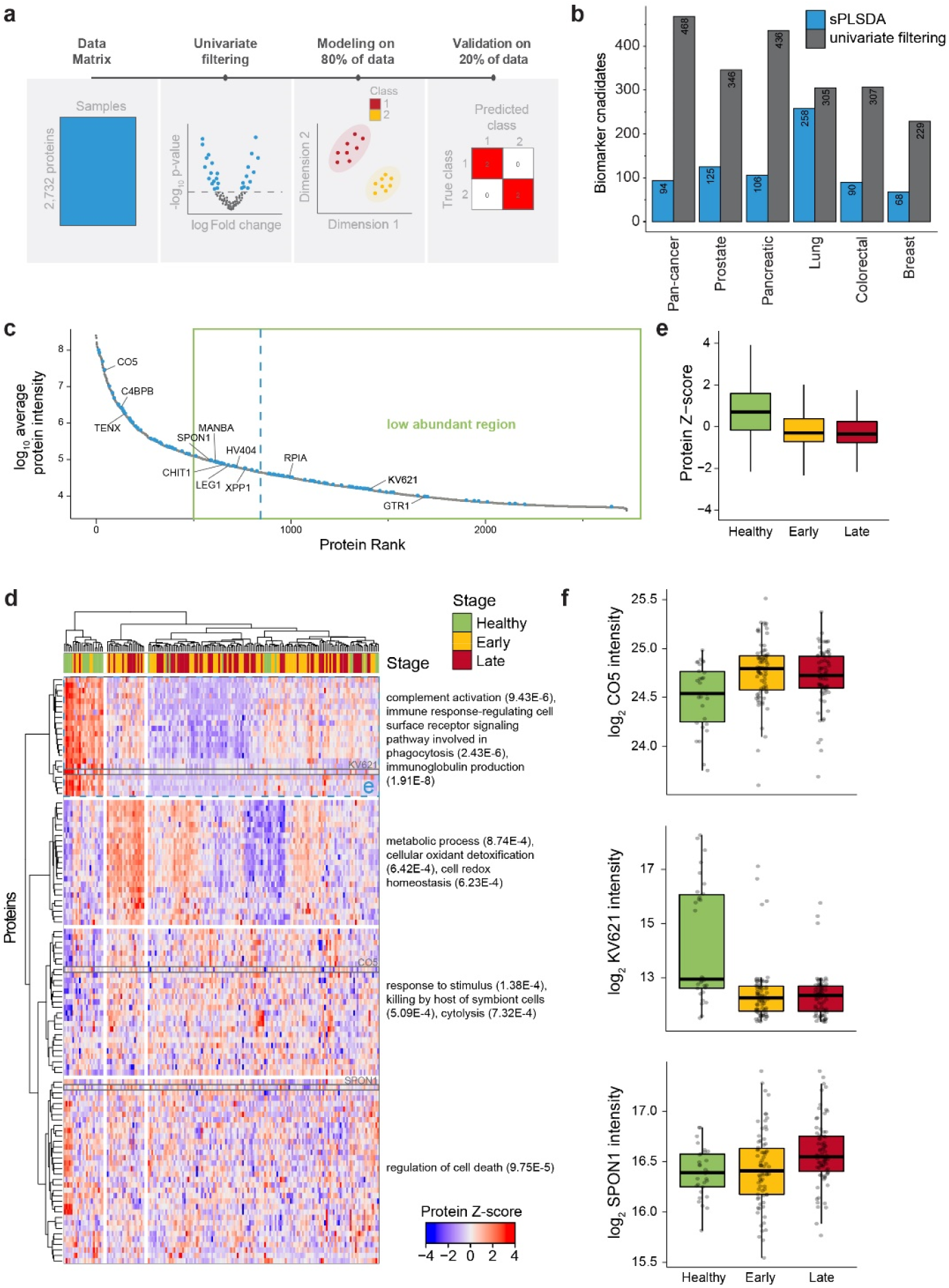
Machine learning-based candidate biomarker discovery. **(a)** Schematic detailing the steps of the post-processing, including univariate testing for filtering, machine learning (sPLSDA) on 80% of the data and classification performance accuracy on the hold-out 20% validation data. **(b)** Overview of the number of biomarker candidates selected by univariate analysis (grey) and machine learning (blue) for healthy, early and late stage, across all cancers and individual cancers. **(c)** Average protein intensity plotted vs. protein abundance rank. The machine learning selected biomarkers candidates for the pan-cancer model are colored in blue (the average is plotted as a blue line) and important contributors are highlighted. The green box depicts the proteome region that is typically below the sensitivity of neat plasma profiling by mass spectrometry. **(d)** Z-score of all machine learning selected candidate biomarkers for the pan-cancer model (n=94) across all measured samples (n=180). Stage calling is overlaid. Both the proteins and the samples were hierarchically clustered. Selected, significantly enriched gene ontology pathways are reported on the right with the p-value in brackets. Proteins highlighted in blue and grey are reported in panels *e* and *f*, respectively. **(e)** Boxplot visualization of the average z-transformed protein intensity for all proteins (n=288) in the cluster highlighted in blue in panel *d* divided by stage (n=180). Thick lines indicate medians, boxes indicate the 25% and 75% quartiles, and whiskers extend between the median and ± (1.58 × interquartile range). **(f)** Boxplot visualization (as in panel *e*) of the log-transformed protein quantities of the three most differentiating proteins based on the machine learning model (SPON1, KV621 and CO5). Each data point represents a sample (n=180).

### Overall changes within and across cancer types

Next, we applied the same analysis strategy using the matched healthy controls to each of the five solid tumor types. In the first step, we identified on average 325 significantly altered proteins between healthy, late and early stages (Fig. 3B and 4A and Supplementary Table 3). With 436 significantly altered proteins (83% reduction in features), prostate cancer had the highest number of differentially abundant proteins, while breast cancer had the fewest with 229 (92% reduction). Interestingly, only a few proteins were shared among cancers (Supplementary Fig. 4A). Pancreatic and prostate had the most with 190 overlapping proteins, while breast and pancreas had the least at 37 (Supplementary Fig. 4A). Seven candidate proteins were consistently selected as differentially abundant across all cancers: the complement activation protein C4b-binding protein beta chain (C4BPB), the immunoglobulin component Immunoglobulin heavy variable 4-4 (HV404), the T-cell apoptosis inducer Galectin-1 (LEG1), the degrader of the inflammation promoting bradykinin peptide Xaa-Pro aminopeptidase 1 (XPP1), the solute carrier family 2 facilitated glucose transporter member 1 (GTR1), the glycan metabolism beta-mannosidase enzyme (MANBA) and the suggested growth inducer of epithelial tumors Tenascin-X (TENX, Fig. 4B and Supplementary Fig. 4A and B). These candidates have rather decreasing (HV404, XPP1, MANBA, TENX) or increasing (LEG1, C4BPB) trends in a cancer agnostic manner, with the exception of GTR1, which strongly increases in late stage breast cancer while decreasing in the other types (Fig. 4C). Interestingly, this small set of proteins separated healthy from the cancer stages samples quite well (p-value = 1.9e-8, Fig. 4B). Fitting a sPLSDA model with 80% of the data overall decreased the number of candidates to less than 5% of the total measured proteins. It led to an average of 129 candidates, making biological interpretation and follow up more feasible (Fig. 3B and 4A and Supplementary Table 4). The relative decrease to the input data was highly cancer dependent, from an almost 76% reduction in pancreatic cancer to only a 15% reduction in lung cancer. The number of overlapping proteins across models was minimal, likely due to the reductionist approach of sPLSDA and cancer type-specific mechanisms, with no proteins being selected for all models (Supplementary Fig. 4C). Still, TAGL and MANBA were selected in all but breast cancer models, and GTR1 and LEG10 in all but the pan-cancer and breast cancer models (Fig. 4C and Supplementary Fig. 4B).

**Fig. 4.**
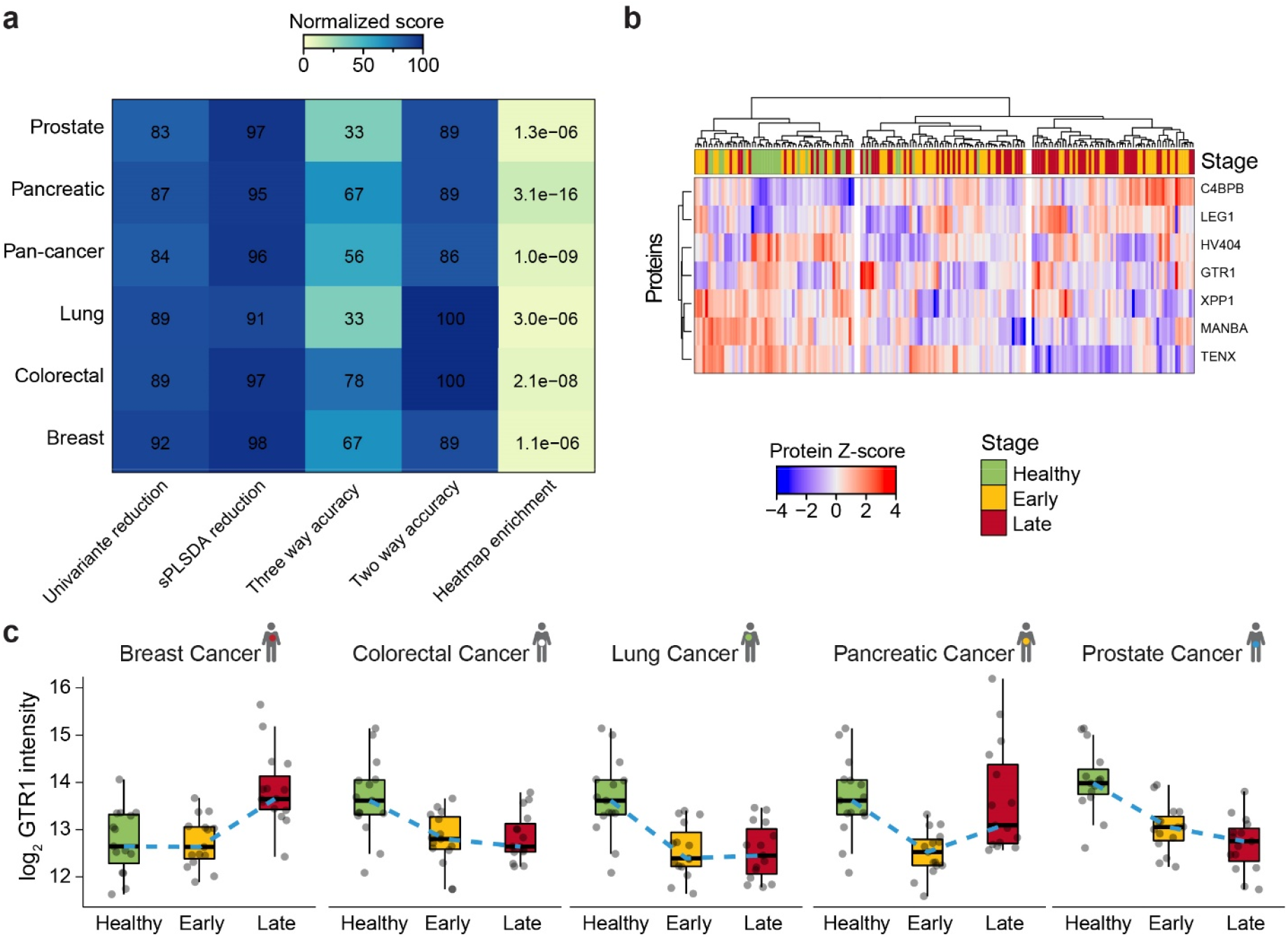
Classification accuracy of the five cancer types. **(a)** Overview of the data analysis per cancer and combined (pan-cancer) as a normalized score. Percentage reduction upon univariate filtering and sPLSDA on 80% of the dataset along with percentage accuracy as measured on the 20% holdout samples as a three-way (healthy, early and late stage) and two-way (cancer and healthy) classification and p-value of enrichment based on the heatmap clustering (Manhattan distance, Ward clustering). **(b)** Z-score of the seven candidate proteins consistently selected across all cancers (by univariate analysis, n=180). Stage calling is overlaid. Both the proteins and the samples were hierarchically clustered. **(c)** Boxplot visualization of log-transformed GTR1 quantities across stage and cancer type. The healthy samples were matched to the respective cancer samples. Thick lines indicate medians, boxes indicate the 25% and 75% quartiles, whiskers extend between the median and ± (1.58 × interquartile range) and each data point represents a sample (n=180). The dashed blue line connects the median values across stages.

In summary, the model classification performance measured on the 20% validation set ranged between 33.3% in lung and prostate cancer to 77.8% in colorectal cancer when all three groups were considered and between 86.1% for the pan-cancer model and 100% for lung and colorectal cancer when healthy and overall disease status were considered (Fig. 4A, Supplementary Table 2). While for the early/late-stage differentiation 2 of the 6 models were close to random performance, the disease status was easier to predict, especially if the cancer type is known, as the pan-cancer model performed the worst with 86% accuracy. Interestingly, high model performance was not always associated with high separation efficiency using PCA or distance analysis and vice versa (Fig. 4A). This is especially apparent in the case of pancreatic and colorectal cancer. While colorectal performs the best on the validation set, especially in the differentiation of healthy/disease, pancreatic cancer leads to the best separation by hierarchical clustering on all three groups (p-value = 3.1e-16). In a nutshell, in contrast to the “one-fits-all” approach, the cancer-specific models performed better. In some cases, the classification accuracy of the derived models was good, demonstrating the benefit of deep profiling of the plasma proteome.

### Disease state separation in colorectal cancer

In colorectal cancer (CRC), we identified 307 proteins significantly altered between healthy, early, and late stages (Supplementary Fig. 5A). The sPLSDA model further reduced these candidate proteins to 90, and both hierarchical clustering and PCA analysis led to efficient separation of healthy subjects from patients regardless of tumor staging (p-value = 2.1e-8, Fig. 5A and Supplementary. Fig 5B). Multiple biological GO enrichments in the candidates could be dissected, for example, response to leptin and regulation of proteolysis increased in cancer (including STAT3 and Transgelin (TAGL)). In contrast, negative regulation of cell-cell adhesion, leukocyte homeostasis and response to hydrogen peroxide decreased (including CD47, Fig. 5A and B). TAGL (importance = 1.00), STAT3 (importance = 0.65) and CD47 (importance = 0.57) were the three most predictive proteins from the sPLSDA model and showed interesting patterns (Fig. 5B and Supplementary Fig. 5C). While CD47 and STAT3 showed strong heterogeneity in late stage colorectal cancer, TAGL was highly expressed in early and late stage colorectal cancer (Fig. 5B). The selected 90 proteins were distributed across the entire intensity range of measured proteins, with more than 80% of the selected proteins (including the most important 3) being beyond the 500 protein mark representing the usual range of proteins detected in neat plasma (Supplementary Fig. 5D). Furthermore, at 78%, the model had the best overall classification accuracy among all tested malignancies on the validation set (Fig. 5C). As no misclassification for healthy subjects was observed, the panel of identified candidate proteins could be helpful for early CRC diagnosis. In summary, despite the small sample set, deep profiling of the human plasma enabled the partial classification of diseased patients based on a panel of 90 proteins that span a large dynamic range while providing an unbiased glimpse into the biological processes at the base of colorectal cancer.

**Fig. 5.**
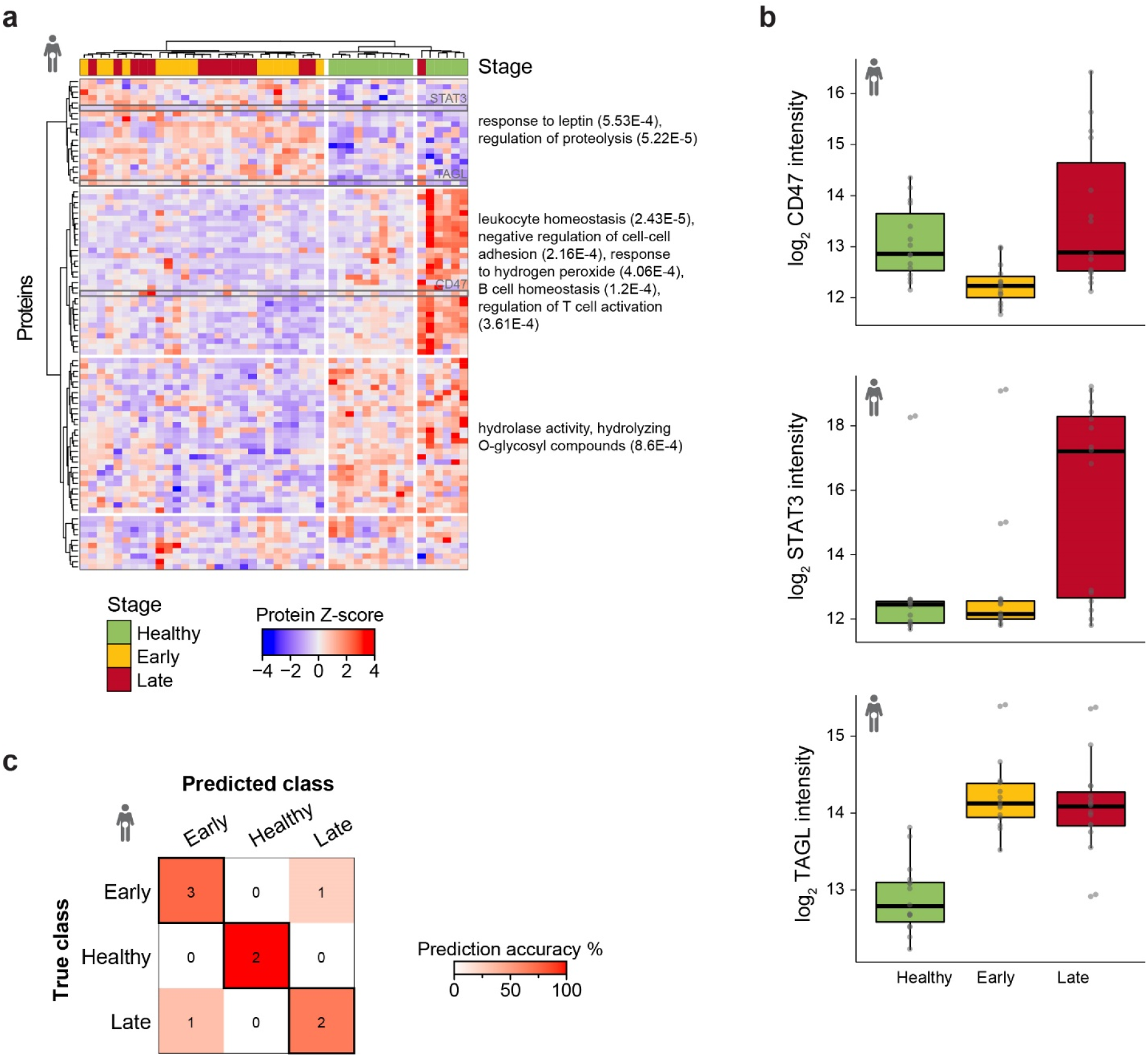
Colorectal cancer biomarker candidates predict disease status. **(a)** Z-score of all machine learning selected candidate biomarkers for the colorectal cancer model (n=90) across the matched colorectal sample set (n=45). Stage calling is overlaid. Both the proteins and the samples were hierarchically clustered. Selected, significantly enriched gene ontology pathways are reported on the right with the p-value in brackets. Proteins highlighted in grey are reported in panel *b*. **(b)** Boxplot visualization of log-transformed CD47, STAT3 and TAGL quantities divided by stage for the colorectal cancer set. Thick lines indicate medians, boxes indicate the 25% and 75% quartiles, whiskers extend between the median and ± (1.58 × interquartile range) and each data point represents a sample (n=45). **(c)** Overview of the classification accuracy of the machine learning models for the colorectal cancer validation set (n=9). Correct classifications are represented in the highlighted boxes.

### Stage separation in pancreatic cancer

In the pancreatic cancer set, 436 proteins were significantly altered between healthy, early and late stages (Supplementary Fig. 6A). The sPLSDA modeling selected 106 proteins, which efficiently separated the three classes in both hierarchical clustering and PCA analyses (p-value = 3.1e-16, Fig. 6A and B). The separation was driven primarily by CD9 (importance = 0.37), TENX (importance = 0.32) and Di-N-acetylchitobiase (DIAC, importance = 0.28), with both TENX and DIAC showing a downregulation with disease progression and CD9 a stronger upregulation in early than late stage pancreatic cancer (Fig. 6C and Supplementary Fig. 6B). CD9 levels correlated most strongly with endocytosis related protein Dynamin-1 (DYN1), Heat shock protein beta-1 (HSPB1), Platelet glycoprotein 4 (CD36) and a profibrotic matricellular protein CCN family member 2 (CCN2). The unsupervised clustering of the candidate proteins resulted in interesting patterns (Fig. 6A). In early stage pancreatic cancer, proteins involved in the regulation of peptide secretion, cell communication and chemokine production are overall downregulated (including LEG10, which is essential for suppressive function of CD25 positive regulatory T-cells^48,49^ (Supplementary Fig. 6C), while proteins involved in negative regulation of apoptotic process and receptor internalization (including Proto-oncogene tyrosine-protein kinase Src (SRC) and CD9, Fig. 6C and Supplementary Fig. 6C) are upregulated. In late stage pancreatic cancer, cellular oxidant detoxification and oxygen transport, including Hemoglobin subunit gamma-1 (HBG1), are upregulated (Supplementary Fig. 6C). Of the 125 biomarker candidates selected, 65% were in the low abundance range (Supplementary Fig. 6D). In the validation set, the model had an accuracy of 66.7%, with two out of nine observations incorrectly assigned to the healthy group instead of the early stage cancer (Fig. 6D). On the whole, deep profiling of human plasma enabled clustering of diseased patients based on disease stage and feature reduction makes biological patterns related to disease progression emerge.

**Fig 6:**
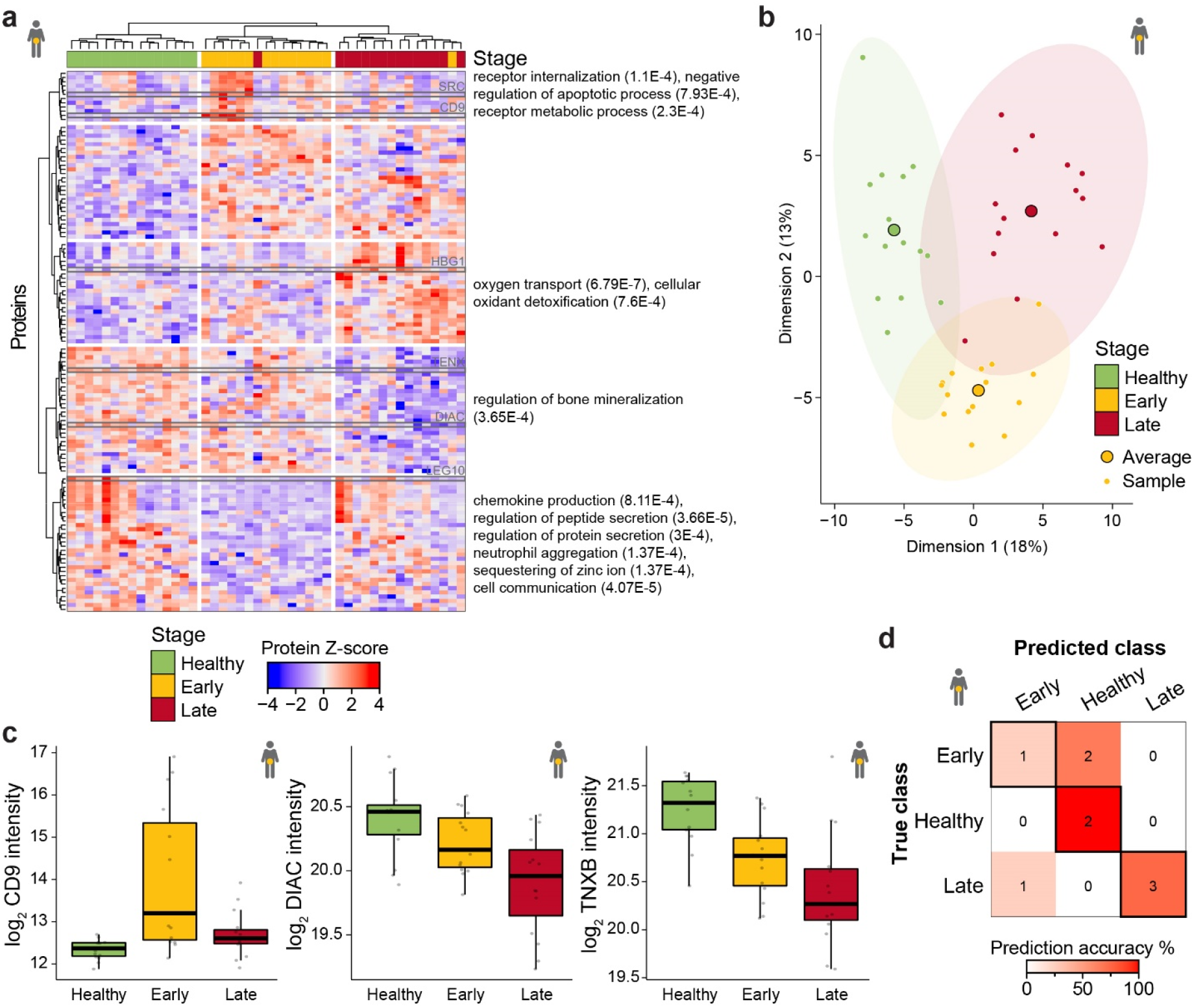
Pancreatic cancer biomarker candidates predict disease stage. **(a)** Z-score of all machine learning selected candidate biomarkers for the pancreatic cancer model (n=106) across the matched pancreatic cancer sample set (n=45). Stage calling is overlaid. Both the proteins and the samples were hierarchically clustered. Selected, significantly enriched gene ontology pathways are reported on the right with the p-value in brackets. Proteins highlighted in grey are reported in panel *c* and the supplementary figure 6. **(b)** Representation of the first two dimensions from the PCA analysis based on candidates identified in the sPLDA model for pancreatic cancer. Small points represent samples and large points the average across the stage. While the first dimension separates healthy from diseased samples and explains 18% of the variance in the data, the second dimension separates early and late stage samples and represents 13% of the variability. Corresponding ellipses represent sample concentration around the mean. **(c)** Boxplot visualization of log-transformed CD9, DIAC and TNXB quantities divided by stage for the pancreatic cancer set. Thick lines indicate medians, boxes indicate the 25% and 75% quartiles, whiskers extend between the median and ± (1.58 × interquartile range) and each data point represents a sample (n=45). **(d)** Overview of the classification accuracy of the machine learning models for the pancreatic cancer validation set (n=9). Correct classifications are represented in the highlighted boxes.

## Discussion

We have developed an automated, robust and parallelizable workflow for deep, large-scale plasma proteome profiling by depletion and sample preparation and by generating deep coverage ion mobility DIA methods. First, we demonstrated substantial improvements upon depletion for identification and quantification using a controlled quantitative plasma experiment. Furthermore, through multistage quality control, we assessed the variance introduced at each step of processing. In summary, the novel plasma discovery workflow enables deep profiling of 10 samples per day per analytical platform to a depth of approximately 2,700 proteins per study for two hours gradients, reaching deep into tissue leakage and signaling molecules while maintaining quantitative accuracy. The protein identifications are expected to increase to about 3,200 cumulatively identified using a four hours gradient FAIMS-DIA acquisitions based on the data from the gradient ramping (Supplementary Fig. 7).

Next, we applied the novel plasma discovery workflow to a cohort containing samples coming from five solid tumors. Data analysis, including machine learning, revealed biomarker candidates and resulted in predictive models. The biomarkers mainly contain proteins from low abundance regions that would have likely been missed by neat plasma profiling, as previously speculated by Geyer et al.^9^.

While separation of healthy from cancer plasma samples was quite accurate for the cancer-specific models (average accuracy 93%), early to late stage differentiation was much more challenging, showing weaker separation (average accuracy 56%). The pan-cancer model performed worse than the cancer-specific models, indicating that “one-fits-all” biomarkers are generally harder to discover. This is likely because of the considerable heterogeneity across cancer types and could be solved by a larger cohort, more advanced stratification strategy and would likely lead to a larger biomarker panel.

Seven candidate proteins were consistently differentially abundant across all cancers, of which one followed a cancer-type specific behavior. Notably, the previously reported pan-cancer biomarker candidate TENX was reproduced, showing a reduction with disease progression irrespective of cancer type^50^. Overall, our approach showed that deep exploration of the proteome of cancer plasma samples can be realized for biomarker discovery. Larger cohorts and a longitudinal study design, where the same subjects are monitored ideally before disease onset would likely lead to more robust biomarkers.

When focusing on colorectal cancer, 307 proteins were altered between healthy, early and late stages. These include three with a documented role in colorectal cancer development: STAT3^51^, TAGL^52^ and CD47^53^. In addition, gene ontology enrichments based on identified candidates showed response to leptin and regulation of proteolysis increased in cancer. At the same time, there was a negative regulation of cell-cell adhesion, leukocyte homeostasis and response to hydrogen peroxide. Based on the machine learning-assisted biomarker discovery approach, a prediction model based on 90 proteins had the highest predictive classification power with 78% accuracy on the hold-out set.

In pancreatic cancer, 436 proteins were altered between healthy, early and late stages. Of these, seven (GTR1, APOA4, IBP2, CD9, CAB45, OLFM4, BGH3) have previously been suggested as possible pancreatic cancer biomarkers ^54–58^. Machine learning-based modeling selected 106 proteins, which led to an efficient separation using distance measures of healthy, early and late stage samples. The selected proteins showed an average overall prediction accuracy of 67%, with two observations incorrectly assigned to the healthy group instead of the early stage cancer. This separation was primarily driven by the three cancer-related proteins CD9^59^, TENX^50^ and DIAC^55,60^. Further proving the quality of the candidates, the separation was also driven by the recently proposed therapeutic target CNN2^61^ and the prognostic marker GTR1^62^. A study by Jayaraman *et al.* demonstrated that exposure of pancreatic cancer cells to zinc leads to increased protein ubiquitination and enhanced cell death, implicating zinc as a potential therapy in treating pancreatic cancer^63^. We found sequestration of zinc ions as an enriched biological process in pancreatic cancer, specifically downregulated in cancer samples (especially early stage).

Clinical analysis of blood is the most widespread diagnostic procedure in medicine, and blood biomarkers are used to diagnose diseases, categorize patients, and support treatment decisions. The presented approach is well suited for deep, epidemiological biomarker studies in plasma as it reaches deep into tissue leakage area, where information on the health state of distal tissues can be discovered. Furthermore, biomarker sets derived from machine learning biomarker discovery analysis are not optimally suited for a direct transition into a “classical” clinical biomarker, as new multiplexed approaches for clinical assays would be required. Such challenges could potentially be facilitated by DIA or multiple PRM-based assays, which are fully compatible with the presented workflow and could ultimately result in streamlined discovery-to-target driven personalized medicine utilizing only one technology platform^64,65^.

Hence, we envision that the profiling of large cohorts at high proteome depth will strongly support the development of novel biomarkers previously not accessible to large-scale discovery approaches and will lead to the development of biomarker panels that will finally deliver on the promise of non-invasive, preventive cancer screening.

## Acknowledgments

We thank Nigel Beaton for input and proofreading the manuscript.

## Abbreviations

CV: Coefficient of variation
DDA: Data-dependent acquisition
DIA: Data-independent acquisition
FDA: Food and drug administration
LC: Liquid chromatography
MS: Mass spectrometry
PTM: Post-Translational Modification

## Author Contributions

R.B., K.S., M.T. and L.R. designed the project. S.M. supported the experimental design of the research. M.T., D.K., J.M. and S.M. developed the sample preparation and prepared the samples. R.B. designed the acquisition methods and M.T. carried out the measurements. M.T and K.S. performed data analysis. M.T., K.S. and R.B. wrote the paper. L.R. supervised the project. All authors critically revised the manuscript and approved its content.

## Data availability

The MS data, the spectral libraries and the quantitative data tables have been deposited to the ProteomeXchange Consortium via the MassIVE repository^66^ with the dataset identifier The Saved projects from Spectronaut can be viewed with the Spectronaut Viewer (www.biognosys.com/spectronaut-viewer).

## Supplementary Figures

**Suppl. Fig. 1:**
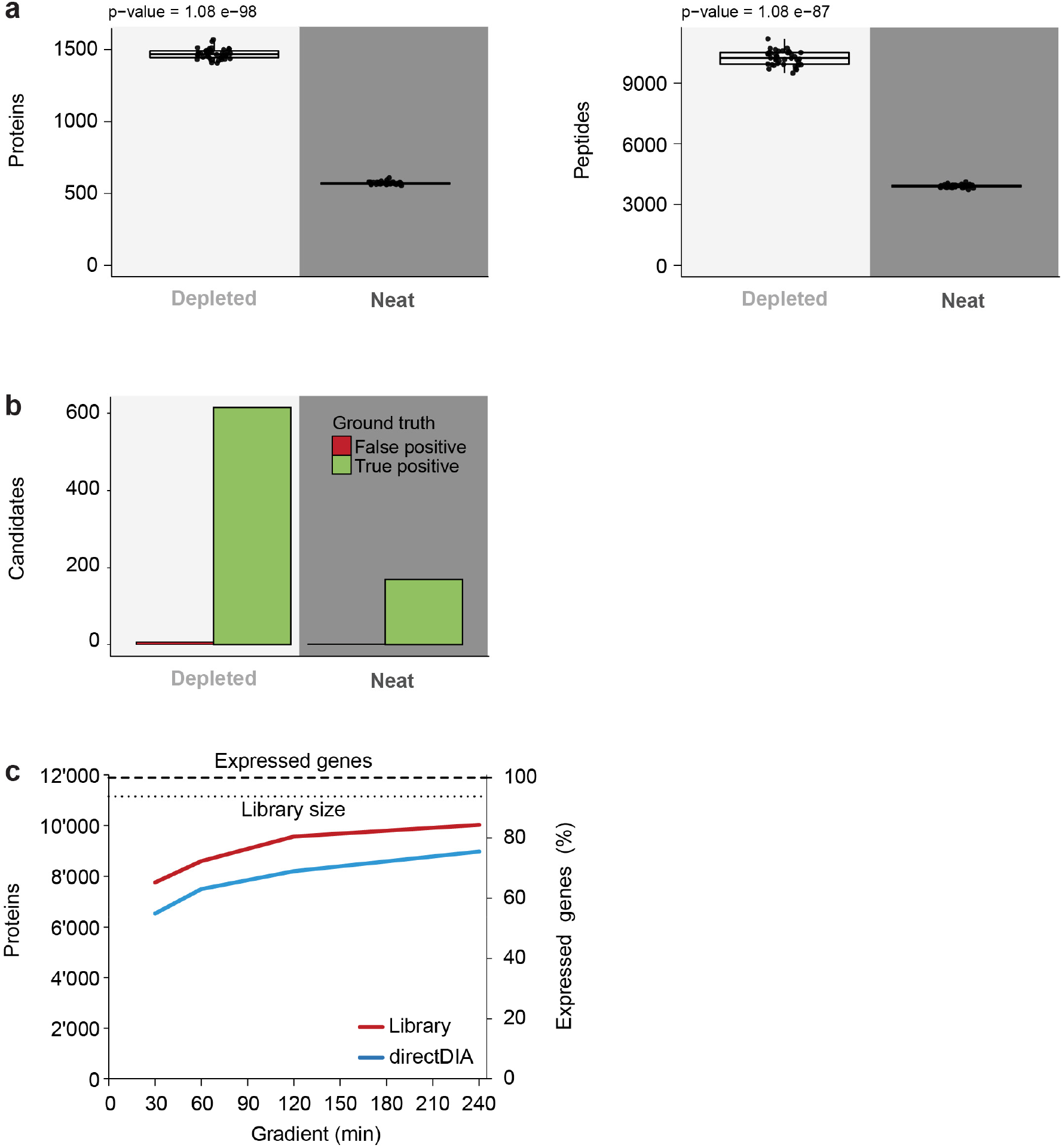
Controlled quantitative experiment with plasma and mass spectrometric performance benchmarking. **(a)** Boxplot visualization of the number of identified protein (protein groups) and peptides (stripped sequences) of the controlled quantitative experiment with neat and depleted plasma. Thick lines indicate medians, boxes indicate the 25% and 75% quantiles, whiskers extend between the median and ± (1.58 × inter-quantile range) and each data point represents a sample (n=80). T-test results are overlaid. **(b)** Representation of the t-test candidates (FDR estimation by the Storey method) divided into true positives and false positives based on the ground truth for the controlled quantitative experiment of both the depleted and neat set. **(c)** Representation of the number of protein identifications from Pierce-HeLa digest using the optimized FAIMS-DIA methods at increasing gradient lengths. The expressed genes number is taken from the human protein atlas and is represented by the thick dashed line (RNAseq data, https://www.proteinatlas.org). The thin dotted line represents the library size.

**Suppl. Fig. 2:**
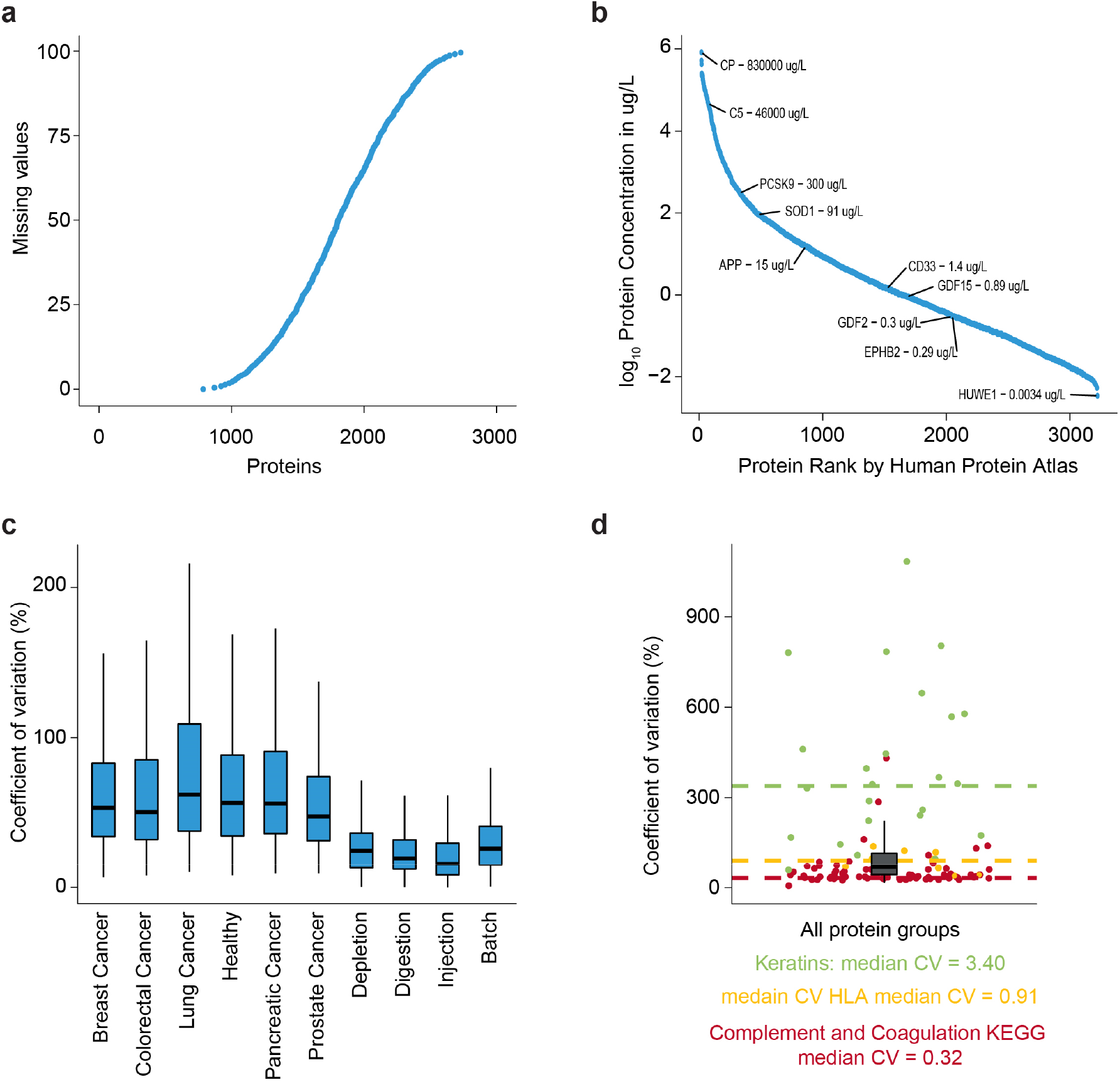
Deep plasma discovery proteomics of five solid cancer types. **(a)** Percentage of missing values in the cohort study plotted against the number of proteins (protein groups) with that value or less. **(b)** Plot of protein rank by the Human Protein Atlas vs. log-transformed reported protein concentration of the identified proteins, spanning 8 orders of magnitude dynamic range as reported in the Human Protein Atlas (3,222 proteins detected in human plasma by mass spectrometry, of which 70% were identified and quantified in this work). Selected proteins were labeled along with the reported concentration. **(c)** Boxplot representation of the coefficient of variation (CV) of the quality control measurements across the processing steps and of the biological variance across cancer types. The CV was calculated on each level: injection (median CV=16%), digestion (CV=19%), depletion (CV=25%), and column (CV=26%). Thick lines indicate medians, boxes indicate the 25% and 75% quantiles, and whiskers extend between the median and ± (1.58 × inter-quantile range). **(d)** Boxplot representation of the biological coefficient of variation across all biological samples (n=180) as in panel *c*. Selected biological pathways are overlaid as points and dashed lines for individual proteins and the median, respectively.

**Suppl. Fig. 3:**
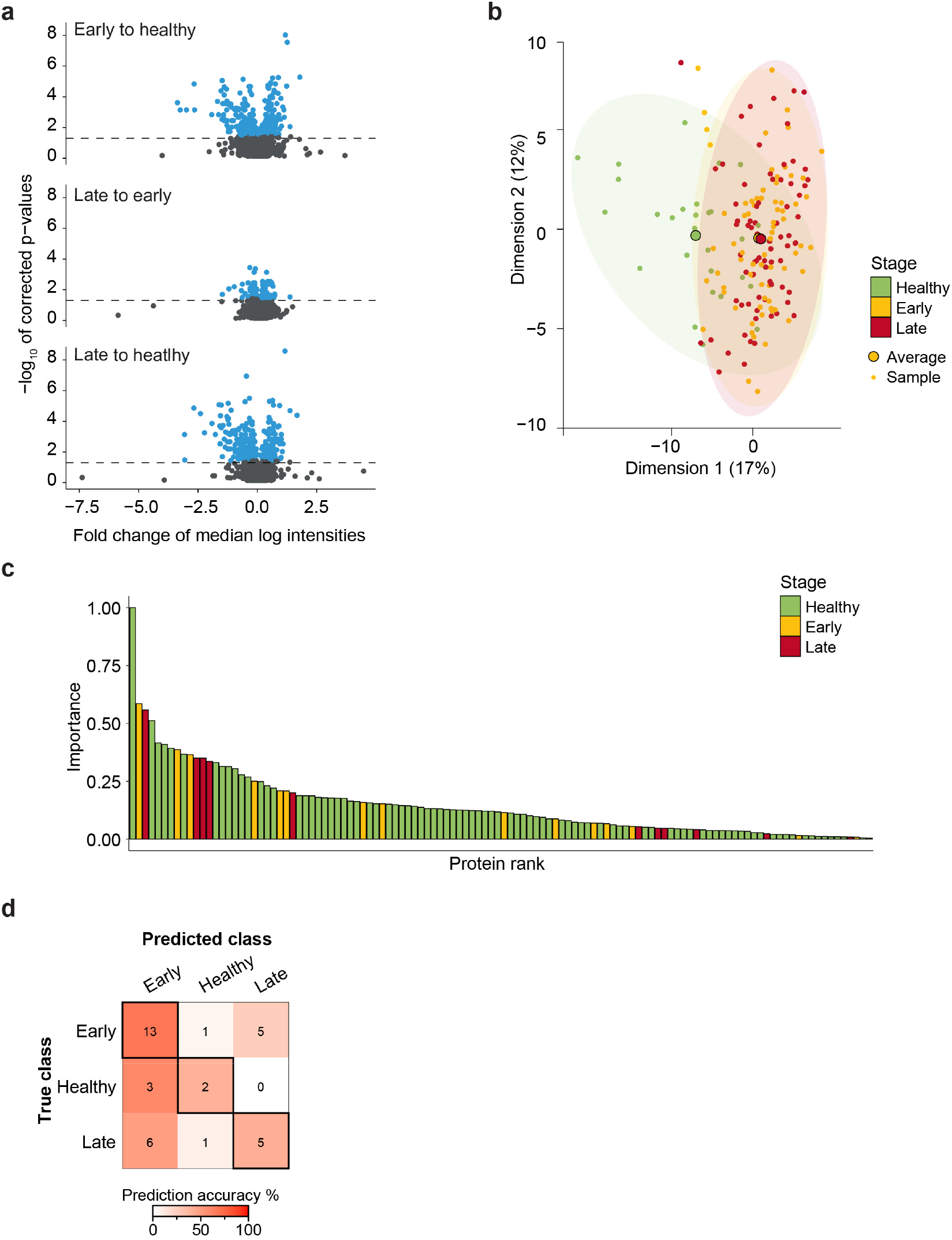
Pan-cancer predictive model based on deeply profiled plasma. **(a)** Log-transformed median fold change vs. –log_10_ p-value for all proteins for the three-way comparisons (heathy, early and late stage) using univariate comparison (Pairwise Wilcoxon Rank Sum Tests) for each protein with all 180 samples. The threshold for protein selection is represented as a dashed line at a p-value 0.05. Proteins with a within-group corrected p-value below 0.05 are depicted in blue. **(b)** Representation of the first two dimensions from the PCA analysis of sPLSDA identified candidates in pan-cancer analysis. Small points represent samples and large points the average across the stage (n=180). The first dimension separates healthy from diseased samples and explains 17% of the variance in the data. Corresponding ellipses represent sample concentration around the mean. **(c)** Representation of the sPLSDA selected biomarker candidates (94 in total) for the pan-cancer model ordered by relative importance and colored by the stage. **(d)** Overview of the classification accuracy of the machine learning model for the pan-cancer validation set (n=36). Correct classifications are represented in the highlighted boxes.

**Suppl. Fig. 4:**
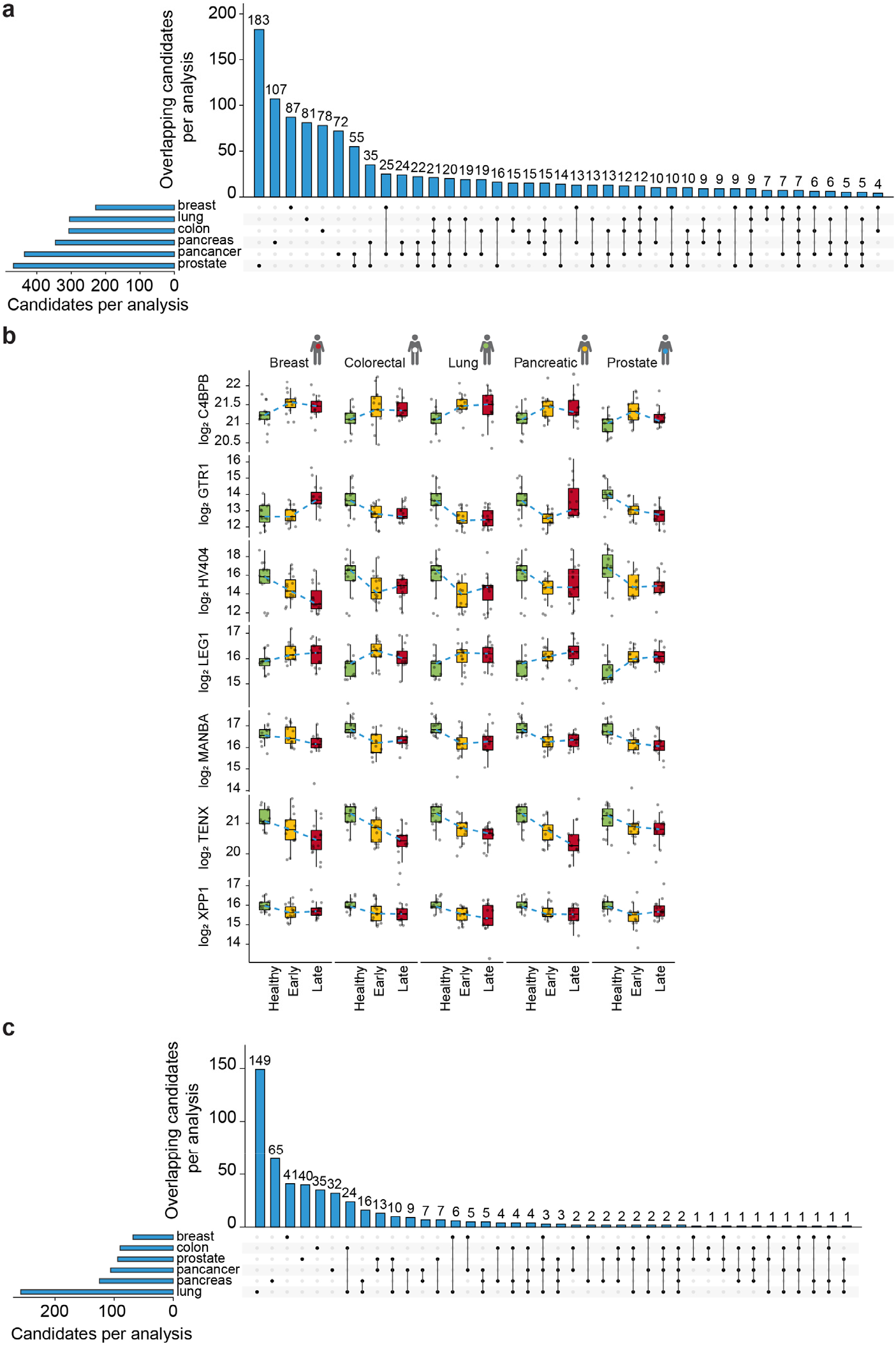
Biomarker candidates within and across the five solid cancers. **(a)** Set plot of proteins coming from the univariate analysis and used downstream for the different cancer models. Blue strips on the left show the number of proteins selected by pairwise comparison. Dots and lines represent subsets. The histogram represents the number of overlapping proteins in each subset. **(b)** Boxplot visualization of log-transformed C4BPB, GTR1, HV404, LEG1, MANBA, TENX and XPP1 quantities divided by stage and cancer type. The healthy samples were matched to the respective cancer type. Thick lines indicate medians, boxes indicate the 25% and 75% quantiles, whiskers extend between the median and ± (1.58 × inter-quantile range), orange lines connect the medians and each data point represents a sample (n=180). The dashed blue line connects the median values across stages. **(c)** As in panel *a* but for the final sPLDA model selections.

**Suppl. Fig. 5:**
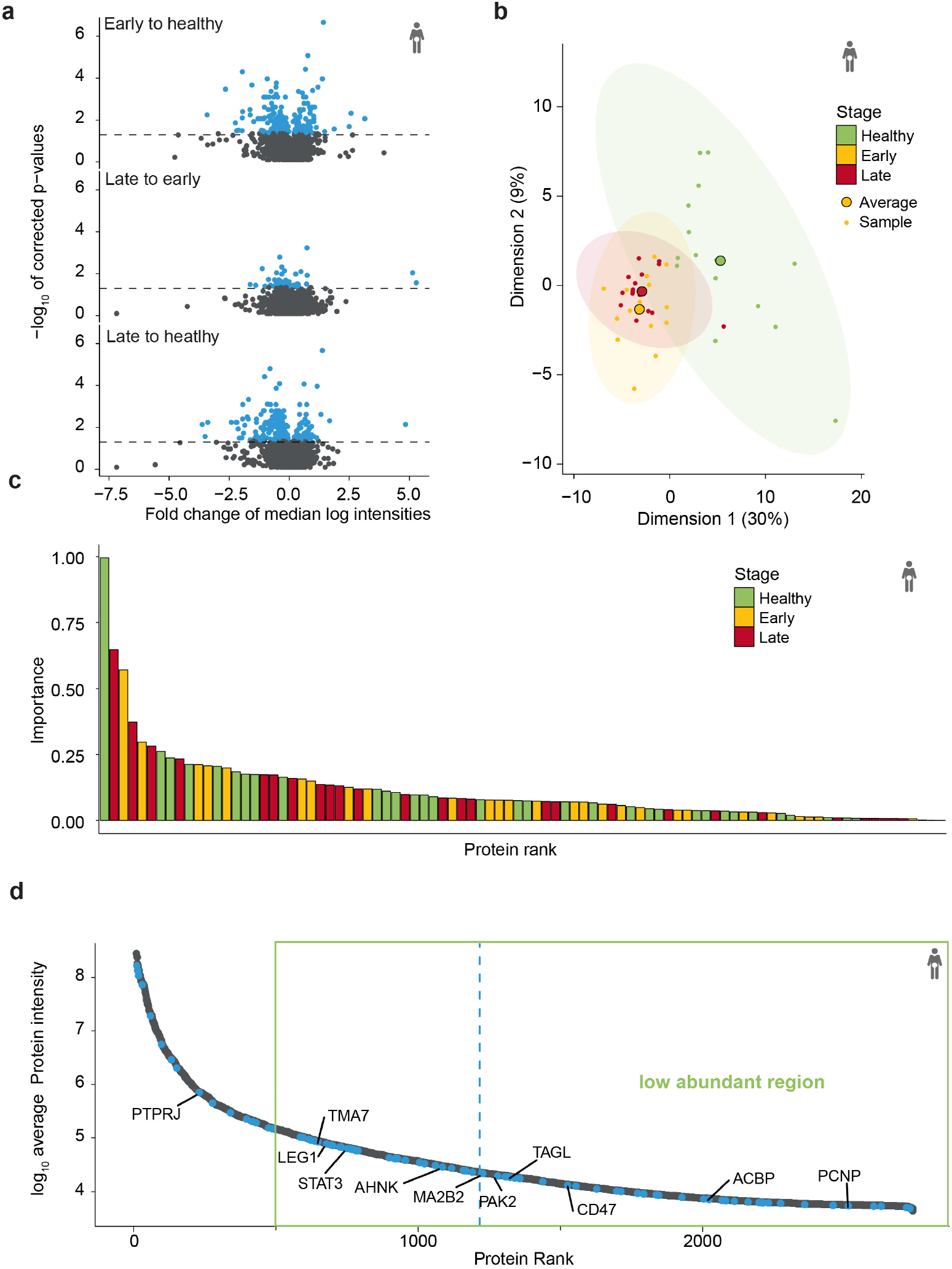
Colorectal cancer analysis. **(a)** Log-transformed median fold change vs. –log_10_ p-value for all proteins for the three-way comparisons (heathy, early and late stage) using univariate comparison (Pairwise Wilcoxon Rank Sum Tests) for the colorectal cancer set (n=45). The threshold for protein selection is represented as a dashed line at a p-value of 0.05. Proteins with a within-group corrected p-value below 0.05 are depicted in blue. **(b)** Representation of the first two dimensions from the PCA analysis of sPLSDA identified candidates in colorectal cancer analysis. Small points represent samples and large points the average across the stage (n=45). The first dimension separates healthy from diseased samples and explains 30% of the variance in the data. Corresponding ellipses represent sample concentration around the mean. **(c)** Representation of the sPLSDA selected biomarker candidates (90 in total) for the colorectal cancer model ordered by absolute importance and colored by the stage. **(d)** Average protein intensity plotted vs. protein abundance rank. The machine learning selected biomarkers candidates for the colorectal cancer model are colored in blue (the average is plotted as a blue line), and important contributors are highlighted. The green box depicts the proteome region that is typically below the sensitivity of native plasma profiling by mass spectrometry.

**Suppl. Fig. 6:**
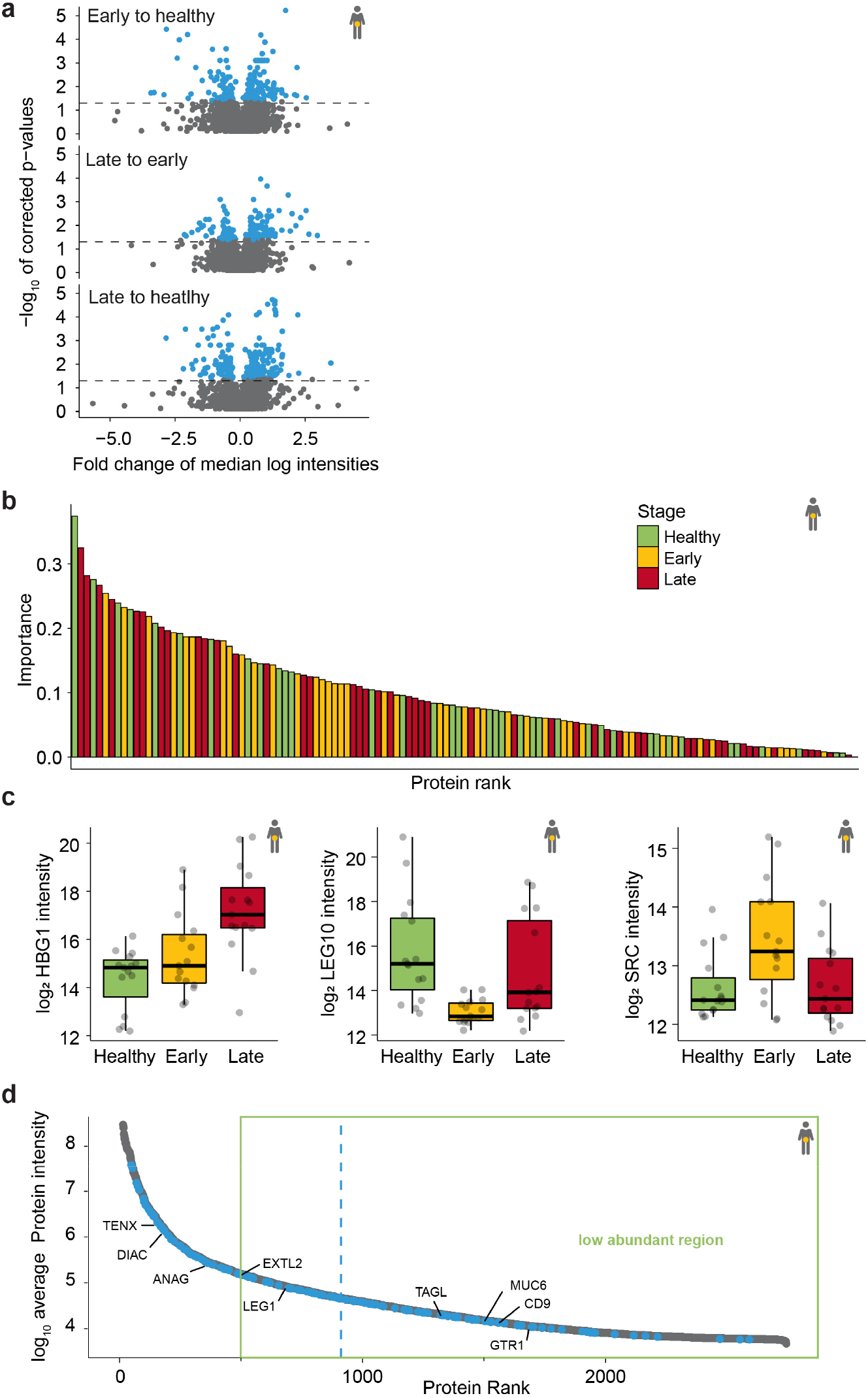
Pancreatic cancer analysis. **(a)** Log-transformed median fold change vs. –log_10_ p-value for all proteins for the three-way comparisons (heathy, early and late stage) using univariate comparison (Pairwise Wilcoxon Rank Sum Tests) for the pancreatic cancer set (n=45). The threshold for protein selection is represented as a dashed line at a p-value of 0.05. Proteins with a within-group corrected p-value below 0.05 are depicted in blue. **(b)** Representation of the sPLSDA selected biomarker candidates for the pancreatic cancer model (106 in total) ordered by absolute importance and colored by the stage. **(c)** Boxplot visualization of selected top candidates log-transformed HBG1, LEG10 and SRC quantities across stages for the pancreatic cancer set. Thick lines indicate medians, boxes indicate the 25% and 75% quantiles, whiskers extend between the median and ± (1.58 × inter-quantile range) and each data point represents a sample (n=45). **(d)** Average protein intensity plotted vs. protein abundance rank. The machine learning selected biomarkers candidates for the pancreatic cancer model are colored in blue (the average is plotted as a blue line) and important contributors are highlighted. The green box depicts the proteome region that is typically below the sensitivity of native plasma profiling by mass spectrometry.

**Suppl. Fig. 7:**
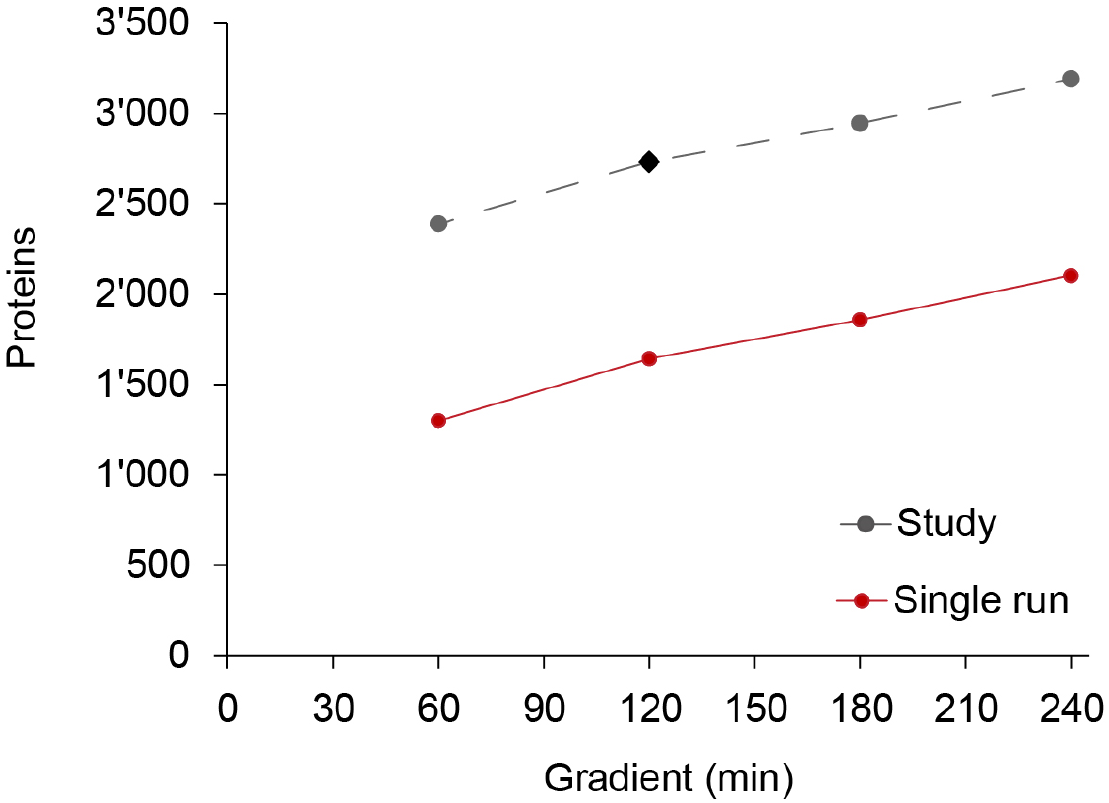
Identifications for a large plasma study in dependence of gradient length. The number of protein groups identified at different gradient lengths for a depleted human plasma pool using a sample specific library (red). The black diamond shows the number of proteins identified in the presented pan-cancer study and the gray dots indicate the extrapolation for different gradient lengths. For the extrapolation, the difference of identifications at 120 minutes of 1,089 proteins was used.

